# Metaproteomics reveals biomarkers of system collapse in a dynamic aquatic ecosystem

**DOI:** 10.1101/040402

**Authors:** Amanda C. Northrop, Rachel Brooks, Aaron M. Ellison, Nicholas J. Gotelli, Bryan A. Ballif

## Abstract

Forecasting and preventing rapid ecosystem state changes is important but hard to achieve without functionally relevant early warning indicators. Here we use metaproteomic analysis to identify protein biomarkers indicating a state change in an aquatic ecosystem resulting from detrital enrichment. In a 14-day field experiment, we used detritus (arthropod prey) to enrich replicate aquatic ecosystems formed in the water-filled pitcher-shaped leaves of the northern pitcher plant, *Sarracenia purpurea*. Shotgun metaproteomics using a translated, custom metagenomic database identified proteins, molecular pathways, and microbial taxa that differentiated control oligotrophic ecosystems that captured only ambient prey from eutrophic ecosystems that were experimentally enriched. The number of microbial taxa was comparable between treatments; however, taxonomic evenness was higher in the oligotrophic controls. Aerobic and facultatively anaerobic bacteria dominated control and enriched ecosystems, respectively. The molecular pathways and taxa identified in the enriched treatments were similar to those found in a wide range of enriched or polluted aquatic ecosystems and are derived from microbial processes that occur at the base of most detrital food webs. We encourage the use of metaproteomic pipelines to identify better early-warning indicators of impending changes from oligotrophic to eutrophic states in aquatic and other detrital-based ecosystems.

## Introduction

Chronic and directional environmental drivers such as nutrient enrichment are causing state changes in many ecosystems (Rabalais et al. 2009, Scheffer 2009). Mitigating or preventing these state changes requires predicting them with sufficient lead-time (Biggs et al. 2009). Current prediction methods rely on the statistical signature of “critical slowing down” (Scheffer et al. 2009)– an increase in the variance or temporal autocorrelation of a state variable (Dakos et al. 2015). However, such indicators usually require long time series of data with frequent sampling of an appropriate state variable (Bestelmeyer et al. 2011, Levin and Mollmann 2015). Even when such data are available, the signature of critical slowing down may not provide enough lead-time for intervention (Biggs et al. 2009, Contamin and Ellison 2009).

In aquatic systems, water quality indicators such as total suspended solids (Hargeby et al. 2007), submersed macrophyte vegetation cover (Dennison et al. 1993, Sondergaard et al. 2010), diatom composition (Pan et al. 1996), and phytoplankton biomass (Carpenter et al. 2008) often are used as state variables. However, whether the change is initiated by top-down or bottom-up forces, the proximate cause of eutrophication in aquatic ecosystems is microbial processes associated with the breakdown of detritus (Chrost and Siuda 2006).

Consequently, a primary reason that it has been difficult to forecast shifts with sufficient lead-time is that changes in monitored variables lag behind the microbial processes that underlie state changes. Here, we describe a molecular pipeline for rapid metagenomic and metaproteomic profiling of microbial composition and function in an experimentally enriched aquatic ecosystem (Fig. 1). This pipeline identified a number of microbial taxa and functional pathways that are common to a wide range of aquatic ecosystems and that have the potential to serve as very early warning indicators of impending state changes in them.

**Fig. 1.**
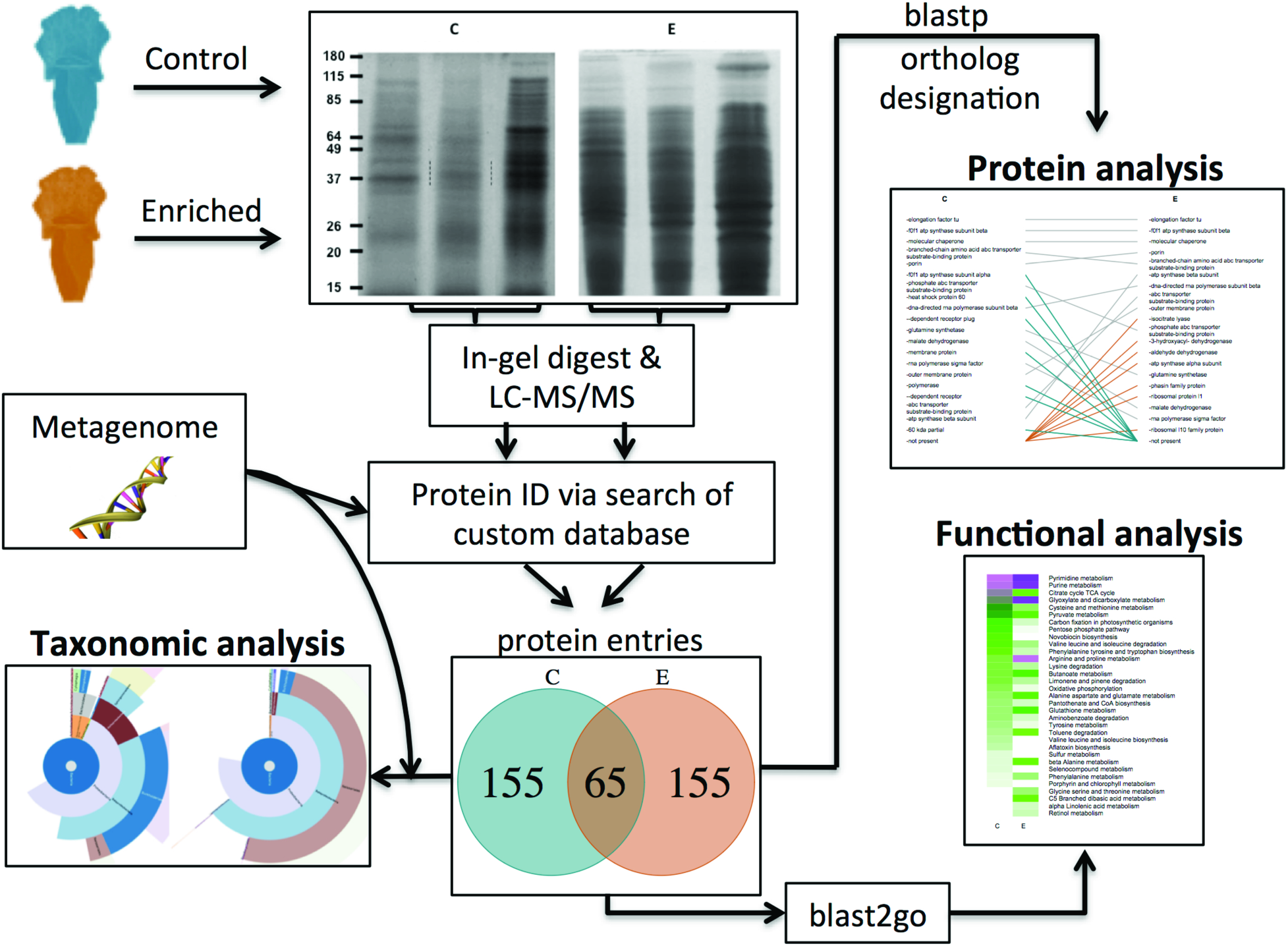
Pipeline for data collection and analysis. Proteins from the microbial communities in experimentally enriched and ambient control pitcher fluid were processed using SDS-PAGE, tryptic digest, LC-MS/MS, and a SEQUEST search of a custom metagenomic database. The composition of microbial communities were determined using a BLAST homology search of metagenomic data associated with identified proteins. Protein identity and annotation was determined via a blastp search to identify orthologs and blast2go.

## Methods

### Study System

The aquatic ecosystem that assembles in the cup-shaped leaves of the northern pitcher plant *Sarracenia purpurea* is an excellent model system for identifying microbial processes driving eutrophication that is triggered by detrital enrichment (Sirota et al. 2013). Each individual leaf functions as an independent replicate ecosystem that can be experimentally enriched and monitored through time in the field or the lab (Srivastava et al. 2004).

Arthropod prey, mostly ants and flies, form the base of a “brown” food web that includes shredding and filter-feeding dipteran larvae and a diverse assemblage of bacteria, primarly Proteobacteria and Bacteroidetes, that decompose and mineralize nearly all of the captured prey biomass (Ellison et al. 2003, Butler et al. 2008, Koopman and Carstens 2011, Gray et al. 2012). Even in the absence of macroinvertebrates, the dominant transfer of nutrients to the plant occurs via microbial activity (Butler et al. 2008).

Under ambient field conditions, prey capture is infrequent (Newell and Nastase 1998), pitcher fluid is clear, and its dissolved oxygen content may exceed 5 mg/L as the plant photosynthesizes in full sunlight (Kneitel and Miller 2002). With detrital loading from excess prey, microbial activity increases, the fluid becomes turbid, and oxygen levels collapse to hypoxic conditions even during daytime photosynthesis (Sirota et al. 2013).

### Enrichment Experiment

In a field experiment conducted in Tom Swamp, a poor fen in central Massachusetts, we randomly assigned 10 pitchers to an ambient control and 10 to a detrital enrichment treatment. Starting June 10, 2011, we selected newly opened pitchers in groups of five for five days until 50 pitchers were selected. One pitcher from each group was randomly assigned to one of five treatments, two of which–control and enriched–are analyzed here. Initial samples of 1.5 ml pitcher fluid were drawn from all pitchers on the first day, and the 1.5 ml of fluid was replaced with 1.5 ml of deionized water. In the detrital enrichment treatment, each pitcher received 1mg/ml/day between 7:00 am and 9:00 am of dried, finely ground wasps (*Dolichovespula maculata*), which has elemental ratios (C:N, N:P:K) similar to those of *Sarracenia’s* natural arthropod prey (Farnsworth and Ellison 2008). Enrichment treatments were applied for 14 consecutive days; pitchers were otherwise unmanipulated.

### Proteomic Analysis

At the end of 14 days, we collected the fluid from each pitcher and filtered it to remove particulates through a 30-μm filter. We centrifuged the filtrate from each replicate separately to concentrate its microbial assemblage, which was then processed using SDS-PAGE (Fig. A1a, Fig. A1b).

Five assemblages from the control pitchers had sufficient microbial biomass for appreciable protein staining, and these were compared to assemblages from six randomly chosen detritus-enriched pitchers, which harbored more microbial biomass. Gel lanes were divided into five regions and the peptides from each gel region were extracted following in-gel tryptic digestion (Fig. A1b). Peptides from each gel region were processed using liquid chromatography tandem mass spectrometry (LC-MS/MS) and were identified using a custom protein database.

We generated this database from a six-frame forward and reverse translation of a metagenomic database that we constructed using the microbial communities of several previously collected pitchers that captured diverse amounts of prey (Fig. A2). The custom database was filtered to only include unique entries (N=118,378) containing at least 100 contiguous amino acids (Appendix A).

### Statistical Analysis

We pooled replicates within treatments and limited our analysis to the 220 most abundant protein identifications in each treatment in order to reduce the false discovery rate of the control and enriched lists to 4.3% and 0%, respectively (Supplement 1). Additionally, the most abundant protein identifications were likely to be important for characterizing ecosystem function and taxonomic coverage.

We conducted a randomization test using R version 3.2.1 (R Project for Statistical Computing) to determine whether there were fewer protein identifications shared between treatments than expected by chance. To determine the taxonomic composition of the microbes that generated these proteins, we used two approaches: 1) a BLAST homology search of the metagenomic sequence data for protein hits, which were then weighted by the number of associated peptides; 2) a search with Unipept (Mesuere et al. 2012, Mesuere et al. 2015) to map tryptic peptides from the samples to the UniprotKB database and retrieve the least common taxonomic ancestor (= most derived shared taxonomic node) associated with each peptide (Appendix A).

To determine whether bacteria in control and enriched ecosystems differed in their O_2_ requirements, we mapped each bacterial species identified in our BLAST search to its O_2_ requirement. We mapped proteins, weighted by the total number of peptides, to metabolic pathways in order to determine functional differences between control and enriched ecosystems (Appendix A).

## Results

Of the 220 most abundant protein identifications, 65 were shared, 155 were unique to control pitchers, and 155 were unique to the enriched treatment (Fig 2a). The randomization test revealed significantly fewer protein hits shared between the treatments than expected by chance (Fig 2b). In both treatments, the top three of the 20 most abundant proteins, as measured by the total number of matched peptides, were the same in the control and enriched treatments. However, the relative abundances of the remaining 17 proteins in this top list differed strongly between treatments, with 7 unique proteins represented only in the enriched treatment and 7 unique proteins represented only in the controls (Fig. 2c).

**Fig. 2.**
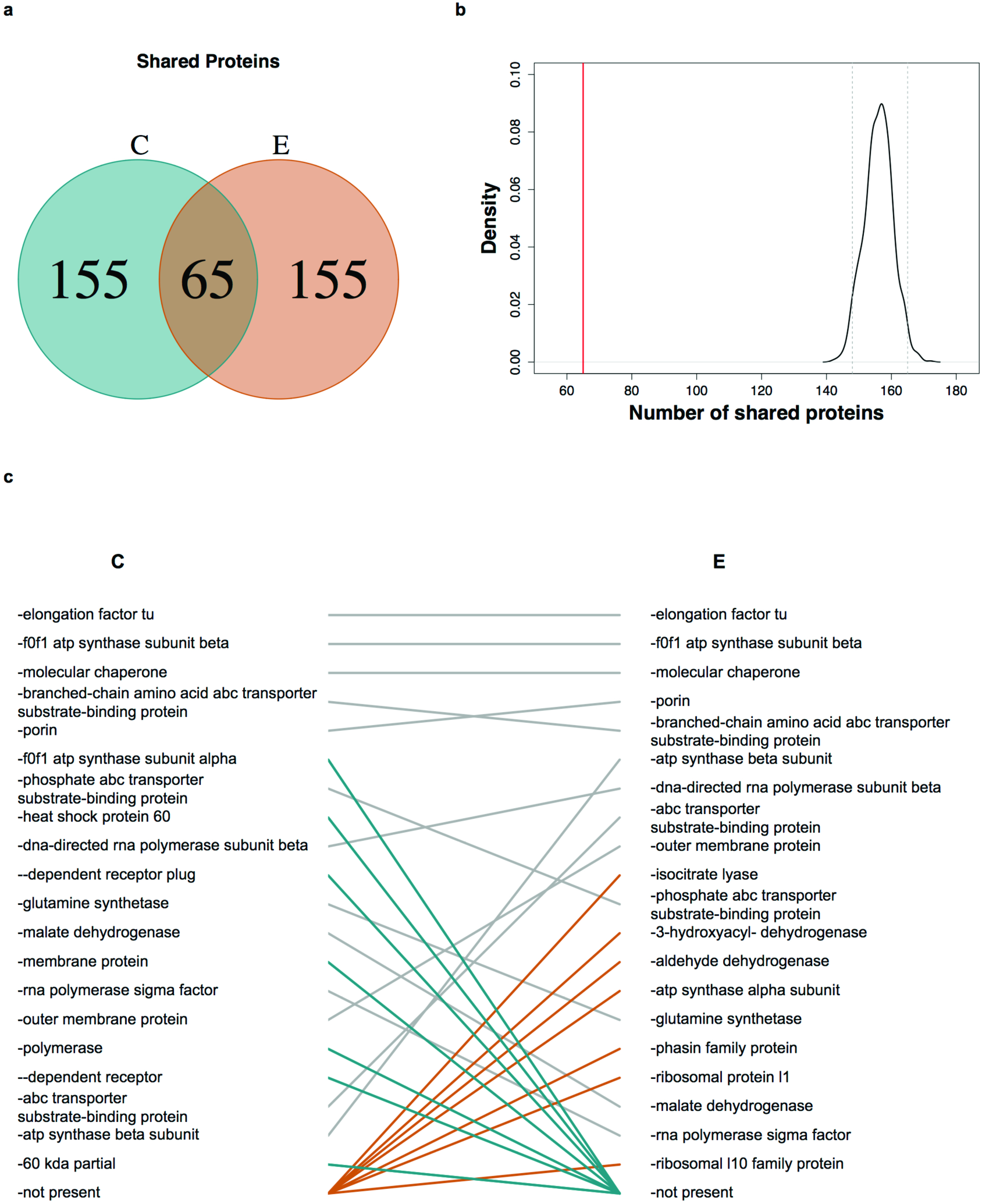
**(a)** Protein hits shared between control and enriched treatments. **(b)** Results of a randomization test in which 220 protein hits were randomly assigned to each treatment and the number of shared protein hits was calculated. Red line indicates the actual shared number of proteins. Grey probability density function indicates the 95% confidence interval for the simulated shared protein hit values. **(c)** Top 20 proteins in rank order for each treatment. Proteins are ranked by the number of total peptides associated with them. Identical proteins in both treatments are connected by lines. Blue lines indicate proteins that were enriched in the control pitchers and brown lines indicate proteins that were enriched in the enriched pitchers.

The most common microbe class in both treatments was Betaproteobacteria, but dominance was higher in enriched (84.4%) versus control (50.3%) treatments (Table A1, Fig. A3). Although both treatments yielded similar numbers of microbe classes (control = 12, enriched = 11), taxonomic evenness was substantially higher in the controls (Hurlbert’s (1971) Probability of an Interspecific Encounter (PIE) = 0.71) than in the enriched treatment (PIE = 0.31). Similar taxonomic profiles were obtained with a search for the least common taxonomic ancestors of the pooled data (Fig. 3a, Fig. 3b). For the unpooled data, taxonomic variability among treatments was greater than variability among replicate ecosystems within treatments (Fig. A4).

**Fig. 3.**
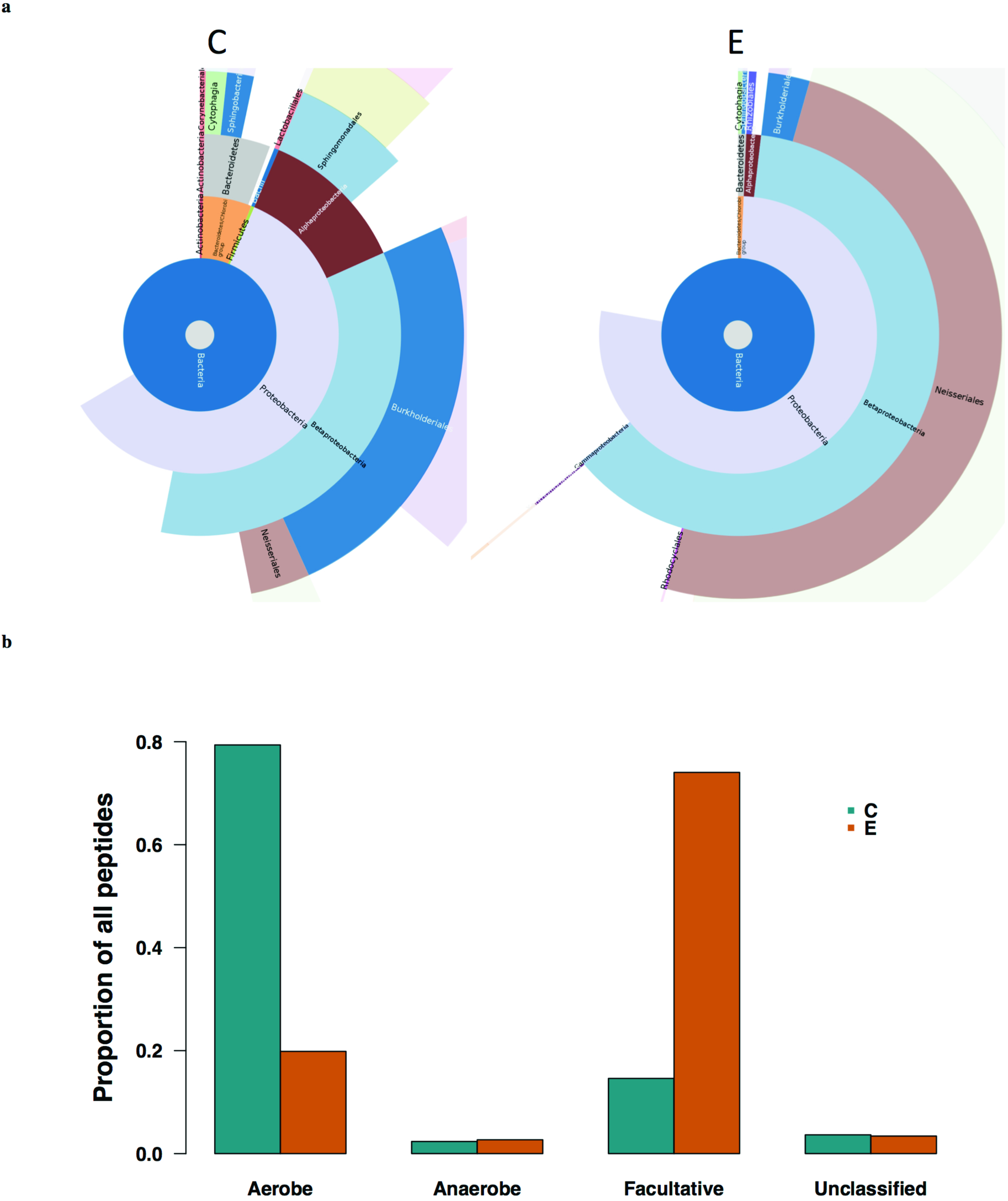
**(a)** Taxonomic composition of bacteria present in control and enriched pitchers visualized using Krona (Ondov et al. 2011). The rings, from the center outward represent Kingdom (dark blue = Bacteria), Phylum (white = Proteobacteria), Class (red = Alphaproteobacteria, light blue = Betaproteobacteria), Order (dark blue = Burkholderiales, rose = Neisseriales, light blue = Sphingomonadales). **(b)** Oxygen requirement classes as a proportion of all peptides for control and enriched pitchers.

Control and enriched treatments differed in function as well as in taxonomic structure. Obligate aerobic bacteria dominated the control pitchers, and facultative anaerobic bacteria dominated the enriched pitchers (Fig. 3c). The difference in oxygen requirement between the two treatments was driven largely by a shift from the obligate aerobe *Variovorax paradoxus* (28.4% of total peptides in the control treatment and 7.2% in the enriched treatment) to the facultative anaerobe *Chromobacterium violaceum* (53.3% of total peptides in the enriched treatment and 6.6% in the control treatment) (Table A2).

Metabolic pathways represented by expressed microbial proteins also differed between control and enriched pitchers. We detected significant differences in metabolic pathways, including those involved in the metabolism of amino acids, carbohydrates, lipids, secondary metabolites, cofactors & vitamins, and terpenoids & polyketides (Fig. 4, Table A3, Fig. A5).

**Fig. 4.**
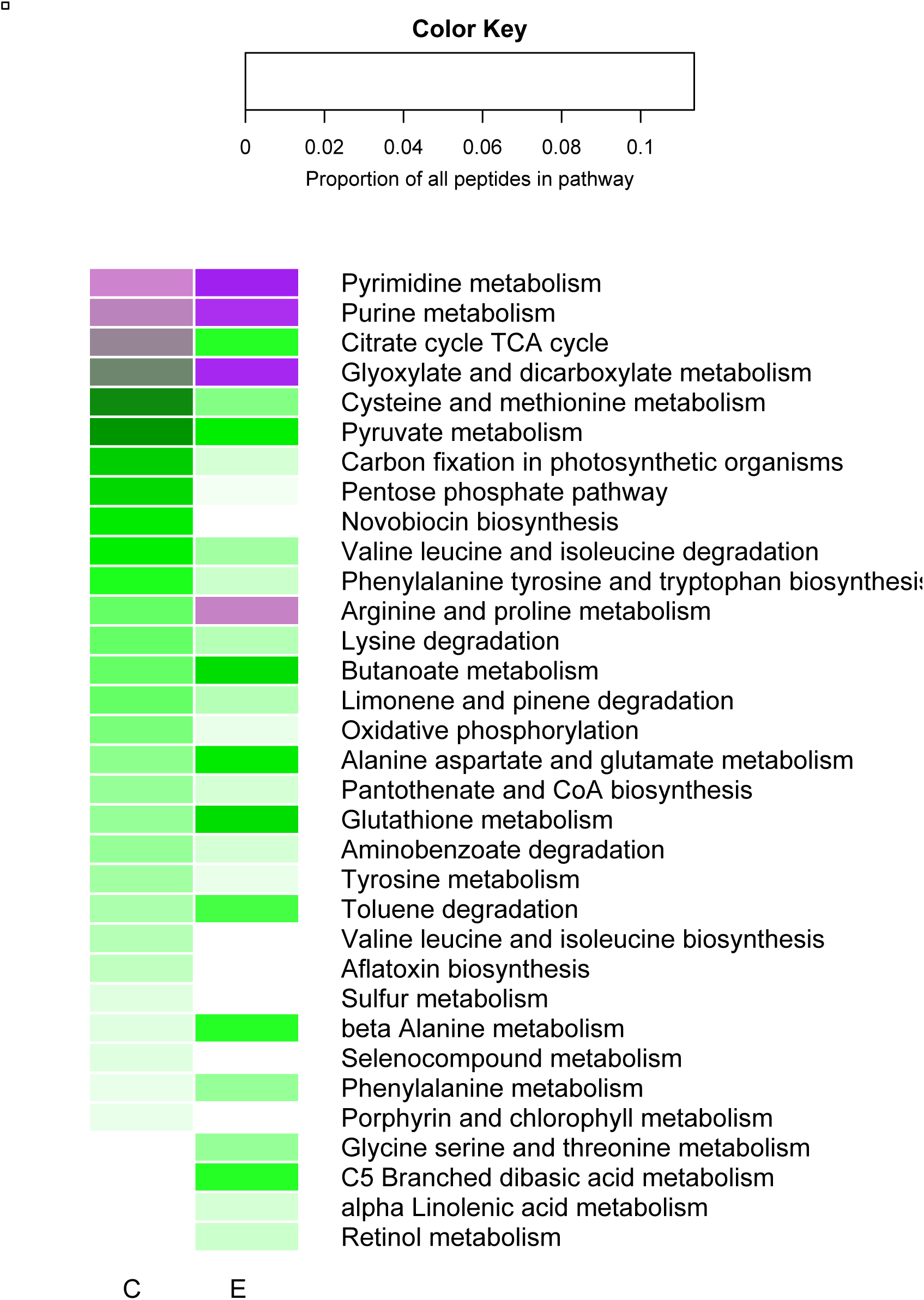
Heat map of the proportional representation of significantly different pathways between control pitchers (C) and enriched pitchers (E).

## Discussion

Changes in land use and climate are likely to exacerbate the causes and symptoms of eutrophication in freshwater aquatic ecosystems. It is therefore becoming increasingly important to predict eutrophic collapse (Jeppesen et al. 2010). Current prediction methods rely on statistical parameters of state variable time series data, but these methods require intensive data collection, are difficult to detect in noisy data sets, and may not provide enough lead-time for successful intervention (Dakos et al. 2015, Bestelmeyer et al. 2011, Contamin and Ellison 2009). Molecular biomarkers, such as proteins produced by the microbial component of aquatic food webs, may be useful early warning signals of impending ecosystem collapse. Screening for such functional indicators of ecosystem change is possible using metaproteomics, which allows us to characterize a wide array of proteins produced by the microbial base of aquatic food webs.

We observed striking differences between oligotrophic (control) and eutrophic (enriched) ecosystems in the molecular function of expressed proteins (Fig. 4) and in the taxonomic composition of the underlying microbial assemblages (Fig. 3). However, these results are not unique or idiosyncratic to this study. The molecular pathways and taxa we identified in experimentally eutrophied *Sarracenia* ecosystems were similar to those found in several other enriched or polluted aquatic ecosystems (Haller et al. 2011, Tang et al. 2009, Debroas et al. 2009).

Patterns of lowered bacterial diversity and increases in Betaproteobacteria abundance displayed by enriched pitchers have been found in other enriched aquatic ecosystems, including contaminated sediments (Haller et al. 2011) and eutrophic lakes (Tang et al. 2009). Several pathways identified in the control and enriched pitchers were common to pathways identified in a mesotrophic lake, including purine metabolism, glyoxylate and dicarboxylate metabolism, and butanoate metabolism (Debroas et al. 2009). Differences in metabolic pathways between treatments may reflect differences in both O_2_ and resource availability that are associated with enrichment.

Differences between control and enriched pitchers suggest that identification and quantitation of metaproteomes at the ecosystem level will reveal candidate biomarkers that can be used as very early warning indicators of state changes in aquatic and other detritus-based ecosystems. In the future, these biomarkers could be used in a Star-Trek "tricorder" for ecologists and citizen scientists to quickly monitor water quality and ecosystem health based on the protein profile of living ecosystems.

## Acknowledgments

This work was funded by the National Science Foundation (grant numbers 1144055 and 1144056). Proteomic analysis was funded by the Vermont Genetics Network through U. S. National Institutes of Health Grant 8P20GM103449 from the INBRE program of the NIGMS. The authors thank Hailee Tenander for assisting with the preparation of samples for mass spectrometry analysis.

## Literature Cited

Bestelmeyer, B. T., A. M. Ellison, W. R. Fraser, K. B. Gorman, S. J. Holbrook, C. M. Laney, M. D. Ohman, D. P. C. Peters, F. C. Pillsbury, A. Rassweiler, R. J. Schmitt, and S. Sharma. 2011. Analysis of abrupt transitions in ecological systems. Ecosphere 2.

Biggs, R., S. R. Carpenter, and W. A. Brock. 2009. Turning back from the brink: Detecting an impending regime shift in time to avert it. Proceedings of the National Academy of Sciences of the United States of America 106:826–831.

Butler, J. L., N. J. Gotelli, and A. M. Ellison. 2008. Linking the brown and green: Nutrient transformation and fate in the Sarracenia microecosystem. Ecology 89:898–904.

Carpenter, S. R., W. A. Brock, J. J. Cole, J. F. Kitchell, and M. L. Pace. 2008. Leading indicators of trophic cascades. Ecology Letters 11:128–138.

Chrost, R. J. and W. Siuda. 2006. Microbial production, utilization, and enzymatic degradation of organic matter in the upper trophogenic layer in the pelagial zone of lakes along a eutrophication gradient. Limnology and Oceanography 51:749–762.

Contamin, R., and A. M. Ellison. 2009. Indicators of regime shifts in ecological systems: What do we need to know and when do we need to know it? Ecological Applications 19:799–816.

Dakos, V., S. R. Carpenter, E. H. van Nes, and M. Scheffer. 2015. Resilience indicators: prospects and limitations for early warnings of regime shifts. Philosophical Transactions of the Royal Society B-Biological Sciences 370.

Debroas, D., J. F. Humbert, F. Enault, G. Bronner, M. Faubladier, and E. Cornillot. 2009. Metagenomic approach studying the taxonomic and functional diversity of the bacterial community in a mesotrophic lake (Lac du Bourget - France). Environmental Microbiology 11:2412–2424.

Dennison, W. C., R. J. Orth, K. A. Moore, J. C. Stevenson, V. Carter, S. Kollar, P. W. Bergstrom, and R. A. Batiuk. 1993. Assessing water-quality with submersed aquatic vegetation. Bioscience 43:86–94.

Ellison, A. M., N. J. Gotelli, J. S. Brewer, D. L. Cochran-Stafira, J. M. Kneitel, T. E. Miller, A. C. Worley, and R. Zamora. 2003. The evolutionary ecology of carnivorous plants. Advances in Ecological Research, Vol 33 33:1–74.

Farnsworth, E. J. and A. M. Ellison. 2008. Prey availability directly affects physiology, growth, nutrient allocation and scaling relationships among leaf traits in 10 carnivorous plant species. Journal of Ecology 96:213–221.

Gray, S. M., D. M. Akob, S. J. Green, and J. E. Kostka. 2012. The bacterial composition within the Sarracenia purpurea model system: Local scale differences and the relationship with the other members of the food web. Plos One 7.

Haller, L., M. Tonolla, J. Zopfi, R. Peduzzi, W. Wildi, and J. Pote. 2011. Composition of bacterial and archaeal communities in freshwater sediments with different contamination levels (Lake Geneva, Switzerland). Water Research 45:1213–1228.

Hargeby, A., I. Blindow, and G. Andersson. 2007. Long-term patterns of shifts between clear and turbid states in Lake Krankesjon and Lake Takern. Ecosystems 10:29–36.

Hurlbert, S. H. 1971. The nonconcept of species diversity: A critique and alternative parameters. Ecology 52:577–586.

Jeppesen, E., Moss, B., Bennion, H., Carvalho, L., DeMeester, L., Friberg, N., Gessner, M.O., Lauridsen, T.L., May, L., Meerhoff, M., Olafsson, J.S., Soons, M.B., Verhoeven, J.T.A. 2010. Interaction of climate change and eutrophication. Pages 119–151 in M. Kernan, R. Batterbee, and B. Moss, editors. Climate change impacts on freshwater ecosystems. Wiley-Blackwell Oxford, UK.

Kneitel, J. M. and T. E. Miller. 2002. Resource and top-predator regulation in the pitcher plant (Sarracenia purpurea) inquiline community. Ecology 83:680–688.

Koopman, M. M. and B. C. Carstens. 2011. The microbial phyllogeography of the carnivorous plant Sarracenia alata. Microbial Ecology 61:750–758.

Levin, P. S. and C. Mollmann. 2015. Marine ecosystem regime shifts: challenges and opportunities for ecosystem-based management. Philosophical Transactions of the Royal Society B-Biological Sciences 370.

Mesuere, B., G. Debyser, M. Aerts, B. Devreese, P. Vandamme, and P. Dawyndt. 2015. The Unipept metaproteomics analysis pipeline. Proteomics 15:1437–1442.

Mesuere, B., B. Devreese, G. Debyser, M. Aerts, P. Vandamme, and P. Dawyndt. 2012. Unipept: Typtic peptide-based biodiversity analysis of metaproteome samples. Journal of Proteome Research 11:5773–5780.

Newell, S. J. and A. J. Nastase. 1998. Efficiency of insect capture by Sarracenia purpurea (Sarraceniaceae), the northern pitcher plant. American Journal of Botany 85:88–91.

Ondov, B. D., N. H. Bergman, and A. M. Phillippy. 2011. Interactive metagenomic visualization in a Web browser. Bmc Bioinformatics 12.

Pan, Y. D., R. J. Stevenson, B. H. Hill, A. T. Herlihy, and G. B. Collins. 1996. Using diatoms as indicators of ecological conditions in lotic systems: A regional assessment. Journal of the North American Benthological Society 15:481–495.

Rabalais, N. N., R. E. Turner, R. J. Diaz, and D. Justic. 2009. Global change and eutrophication of coastal waters. Ices Journal of Marine Science 66:1528–1537.

Scheffer, M. 2009. Critical Transitions in Nature and Society. Princeton University Press.

Scheffer, M., J. Bascompte, W. A. Brock, V. Brovkin, S. R. Carpenter, V. Dakos, H. Held, E. H. van Nes, M. Rietkerk, and G. Sugihara. 2009. Early-warning signals for critical transitions. Nature 461:53–59.

Sirota, J., B. Baiser, N. J. Gotelli, and A. M. Ellison. 2013. Organic-matter loading determines regime shifts and alternative states in an aquatic ecosystem. Proceedings of the National Academy of Sciences of the United States of America 110:7742–7747.

Sondergaard, M., L. S. Johansson, T. L. Lauridsen, T. B. Jorgensen, L. Liboriussen, and E. Jeppesen. 2010. Submerged macrophytes as indicators of the ecological quality of lakes. Freshwater Biology 55:893–908.

Srivastava, D. S., J. Kolasa, J. Bengtsson, A. Gonzalez, S. P. Lawler, T. E. Miller, P. Munguia, T. Romanuk, D. C. Schneider, and M. K. Trzcinski. 2004. Are natural microcosms useful model systems for ecology? Trends in Ecology & Evolution 19:379–384.

Tang, X. M., G. Gao, B. Q. Qin, L. P. Zhu, J. Y. Chao, J. J. Wang, and G. J. Yang. 2009. Characterization of bacterial communities associated with organic aggregates in a large, shallow, eutrophic freshwater lake (Lake Taihu, China). Microbial Ecology 58:307–322.

